# Effects of Retatrutide on Learning and Memory in Streptozotocin-Induced Male Diabetic Rats

**DOI:** 10.64898/2026.01.23.701347

**Authors:** Ulya Keskin, Eslem Altin, Melkan Kagan Kara, Burak Tekin, Kerime Nur Cakircoban, Fikriye Yasemin Ozatik, Neziha Senem Ari, Ayse Kocak Sezgin, Emre Gungor

## Abstract

Diabetes mellitus is associated with cognitive impairment and neurodegenerative changes, partly through hyperglycaemia-driven neuroinflammation and disrupted neuronal signalling. Retatrutide, a triple GIP/GLP-1/glucagon receptor agonist, has shown strong metabolic efficacy, but its effects on diabetes-associated cognitive dysfunction remain unclear. The present study investigated whether Retatrutide attenuates learning- and memory-related impairments in a streptozotocin-induced, insulin-deficient diabetic rat model. Male Sprague-Dawley rats were allocated to four groups: control (C), streptozotocin-induced diabetic (STZ), streptozotocin-induced diabetic treated with Retatrutide (STZR), and Retatrutide alone (R). Diabetes was induced with streptozotocin, and spatial learning and memory were assessed using the Morris Water Maze and Passive Avoidance tests. Metabolic parameters were monitored, while hippocampal cytokine levels (IL-1β, TNF-α), BDNF, CREB, and AKT mRNA expression, Tau protein levels, and cortical and hippocampal histopathology were evaluated using biochemical, molecular, and histological methods.

Streptozotocin-induced diabetes produced persistent hyperglycaemia, marked body weight loss, and impaired behavioural performance, particularly prolonged escape latencies in the Morris Water Maze and a selective short-term Passive Avoidance deficit. Retatrutide reduced blood glucose levels but did not prevent diabetes-associated weight loss. In behavioural testing, Retatrutide-treated diabetic rats showed preserved overall Morris Water Maze performance relative to untreated diabetic rats and a limited, task-dependent attenuation of short-term avoidance deficits rather than complete normalisation across all memory measures. These effects were accompanied by a significant reduction in hippocampal TNF-α, a non-significant trend toward lower IL-1β, and partial preservation of cortical and hippocampal cytoarchitecture. Retatrutide alone did not improve behavioural performance beyond control levels, although BDNF and CREB mRNA expression were increased in the non-diabetic Retatrutide group. These findings indicate that Retatrutide is associated with a partial attenuation of streptozotocin-induced behavioural and neuroinflammatory alterations in male rats. The observed effects are consistent with actions extending beyond glycaemic control alone, although direct central exposure of Retatrutide was not established in the present study. Further studies in insulin-resistant and type 2 diabetes-like models are needed to clarify the underlying mechanisms and translational relevance.

## 1. Introduction

Diabetes is a chronic metabolic disease characterised by persistent hyperglycemia, which affects multiple organ systems and leads to various complications [1]. It has become a major global public health concern.

A large-scale meta-analysis including data from 1108 individuals reported that, as of 2022, approximately 828 million adults worldwide were living with diabetes, reflecting a dramatic rise in prevalence over recent decades [2]

Type 2 diabetes causes dysfunction in multiple organs, including the brain and nervous system. Cognitive impairments, particularly those involving learning and memory, are frequently observed as central nervous system complications. Studies have shown that diabetic patients have a 50% higher risk of developing Alzheimer’s disease, vascular dementia, and other neuropsychological disorders compared to non-diabetic individuals [3]. Moreover, cognitive deficits related to diabetes occur not only in older adults but also in adolescents [1]. In this context, developing treatment strategies for cognitive impairments in prediabetic and diabetic adolescents is of great importance.

Glucagon-like peptide-1 (GLP-1) receptor agonists, also known as incretin mimetics, are widely used in the treatment of type 2 diabetes and obesity [4,5]. Glucose-dependent insulinotropic polypeptide (GIP) is another incretin hormone that modulates the secretion of insulin and glucagon (GCG). Preclinical studies have demonstrated that triple receptor agonists (RAs) activating all three receptors (GLP-1, GIP, and GCG) produce more pronounced weight loss than dual agonists targeting only two receptors [6]. Therefore, such agents are expected to become increasingly utilised. Retatrutide is a novel triple hormone agonist that simultaneously activates GIP, GLP-1, and GCG receptors. Both *in vivo* and *in vitro* studies in humans and animals are currently investigating its therapeutic potential for obesity and diabetes [6].

Animal studies have revealed that GLP-1 receptor agonists can cross the blood-brain barrier and distribute within various brain regions. Additionally, GLP-1 is endogenously synthesised in the nucleus of the solitary tract. Numerous preclinical studies have investigated the neuroprotective effects of GLP-1 analogues, demonstrating that they enhance neuronal survival and repair, promote mitochondrial biogenesis, and exert antioxidant and anti-inflammatory effects in the brain [7].

Proinflammatory cytokines also play a crucial role in diabetes-related cognitive decline. Tumour necrosis factor-alpha (TNF-α) released from microglia or perivascular macrophages activates the c-Jun N-terminal kinase (JNK) and inhibitor of κB alpha/nuclear factor κB (IκBα/NF-κB) signalling pathways, thereby amplifying neuroinflammatory responses throughout the brain parenchyma [8]. In diabetic patients, increased circulating levels of inflammatory markers such as C-reactive protein (CRP), interleukin-6 (IL-6), and TNF-α have been associated with poorer cognitive performance [9]. Similarly, elevated IL-1β levels have been observed in elderly patients with mild cognitive impairment compared with healthy controls [10]. Furthermore, a cross-sectional study involving 1712 diabetic patients found that chronic inflammatory diseases, such as rheumatoid arthritis and asthma, also constitute additional risk factors for cognitive decline in diabetes [11].

Brain-derived neurotrophic factor (BDNF) is a neuroprotective growth factor that supports neuronal proliferation, differentiation, and survival [12]. BDNF binds to the tropomyosin receptor kinase B (TrkB) receptor, promoting neuronal survival and plasticity through the activation of key intracellular signalling cascades, including the phosphoinositide 3-kinase/protein kinase B (PI3K/AKT), phospholipase C-γ (PLC-γ), and mitogen-activated protein kinase/extracellular signal-regulated kinase (MAPK/ERK) pathways [13,14]. These signalling events ultimately enhance cAMP response element-binding protein (CREB) activation and regulate gene transcription in neurons. The BDNF/TrkB signalling axis thus plays an essential role in neuronal survival, memory formation, and antidepressant-like effects [15,16].

Streptozotocin (STZ) is a glucosamine-nitrosourea compound that selectively destroys pancreatic β-cells and is widely used to induce experimental diabetes models. Streptozotocin was used as the diabetogenic agent in this study. To avoid terminological ambiguity, the term Streptozotocin is written in full when referring to the compound, whereas STZ and STZR are retained only as experimental group labels. In the present study, diabetes was induced using Streptozotocin at a dose of 60 mg/kg [17,18]. Although this insulin-deficient model primarily resembles type 1 diabetes mellitus (T1DM-like), it provides a robust and widely accepted platform to investigate diabetes-associated central nervous system alterations, particularly hyperglycaemia-driven neuroinflammation and cognitive impairment. Therefore, the present study focused on diabetes-related neurocognitive changes rather than modelling a specific diabetes subtype. In this context, the present study was designed to evaluate whether Retatrutide exerts direct or indirect neuroprotective effects on learning and memory that extend beyond its metabolic actions.

The increasing prevalence of diabetes and obesity, particularly at younger ages, underscores the importance of understanding how antidiabetic drugs influence cognitive function. The present study was designed as an initial proof-of-concept experiment to characterise the behavioural and hippocampal molecular effects of Retatrutide in a well-established drug-induced rat model of diabetic cognitive impairment. As an initial proof-of-concept experiment, the study was restricted to male rats to minimise potential variability related to the oestrous cycle and to preserve comparability with previous Streptozotocin-based studies of diabetes-associated cognitive impairment; however, this design does not permit conclusions about sex-dependent effects. This design was intended to provide mechanistic preclinical evidence under controlled experimental conditions rather than to reproduce the full clinical heterogeneity of human diabetes. To date, no studies have examined the cognitive effects of Retatrutide, a novel triple GLP-1/GIP/GCG receptor agonist. Therefore, this study aimed to investigate the effects of Retatrutide on learning and memory performance in Streptozotocin-induced diabetic male rats.

## 2. Materials and Methods

### 2.1. Ethical Approval

The study was approved by the Kütahya Health Sciences University (KHSU) Animal Experiments Local Ethics Committee (Decision No: 2024.01.06, Date: 30.01.2024). The project was supported by the KHSU Scientific Research Projects Coordination Unit.

### 2.2. Chemicals

Retatrutide was obtained from ALFAGEN Laboratory Supplies Co. (İzmir, Türkiye). Streptozotocin was obtained from certified laboratory-grade sources. Retatrutide was first dissolved in 100% DMSO to prepare a stock solution. Immediately prior to injection, the stock was diluted with saline to obtain a final formulation containing 10% DMSO and 90% saline (v/v). All animals received corresponding vehicle solutions in equal volumes to eliminate solvent-related confounding effects. Streptozotocin was dissolved freshly in cold 0.1 M citrate buffer (pH 4.5) immediately before intraperitoneal injection.

### 2.3. Animals and Housing

A total of 36 male Sprague Dawley rats (10-12 weeks old) were used. Animals were housed at 24 ± 2 °C, 55 ± 15% humidity, under a 12 h light/dark cycle, with ad libitum access to water and standard chow. All experimental procedures complied with national and international guidelines for the care and use of laboratory animals. Experiments were conducted at the KHSU Experimental Animal Breeding Research and Application Centre and at the Department of Medical Pharmacology.

### 2.4. Experimental Design

Rats were allocated into four groups using stratified randomisation based on Barnes Maze performance: Control (C, n = 8); Streptozotocin (STZ, initial n = 9, final n = 5); STZ + Retatrutide (STZR, initial n = 11, final n = 6); and Retatrutide (R, n = 8). Initial group sizes were determined in line with previous Streptozotocin-based behavioural studies, with deliberate oversampling of diabetic groups-particularly STZR– to account for anticipated diabetes-related mortality and exclusion due to severe metabolic or locomotor impairment. For clarity, Streptozotocin is referred to in full throughout the text when describing the diabetogenic compound and its administration, whereas STZ and STZR are used only as group identifiers.

Diabetes was induced in the STZ and STZR groups by a single intraperitoneal injection of Streptozotocin (60 mg/kg). All injections were administered in a total volume of 2 mL per rat (STZ/STZR: Streptozotocin solution; C/R: citrate buffer). Animals with blood glucose levels <250 mg/dL after the initial injection received an additional Streptozotocin dose (30 mg/kg, 2 mL); this was observed in one rat in the STZ group and two rats in the STZR group. However, none of these animals survived to the planned assessment stages and were therefore excluded from all analyses. Accordingly, all reported behavioural and molecular data were derived from animals that received a single Streptozotocin dose (60 mg/kg) and survived to the experimental endpoint.

Before behavioural testing, 3 rats in the STZ group and 5 in the STZR group did not survive; these losses included all animals that received a supplementary Streptozotocin dose. In addition, one rat in the STZ group developed severe clinical deterioration with marked locomotor impairment and was euthanised; this animal was excluded from Morris Water Maze (MWM) and Passive Avoidance (PA) testing, but its tissue samples were retained for molecular and histological analyses. For molecular outcomes, outliers exceeding ±3 standard deviations from the group mean were excluded based on predefined criteria (IL-1β: one C, two R; Tau: one STZR).

Diabetes was confirmed three days after injection, with blood glucose levels ≥250 mg/dL considered diabetic. Retatrutide treatment was initiated immediately thereafter and administered subcutaneously at 0.015 mg/kg once daily for 21 consecutive days using a working solution containing 10% DMSO and 90% saline (v/v), in a fixed volume of 1 mL per rat, at approximately the same time each day to minimise circadian variability. To equalise solvent exposure, citrate buffer was administered to the C and R groups corresponding to Streptozotocin treatment, and 10% DMSO + 90% saline was administered to the C and STZ groups corresponding to Retatrutide treatment, using the same volume and schedule.

Body weight was recorded after Barnes Maze testing (baseline) and immediately before sacrifice. Blood glucose levels were measured on the day of diabetes confirmation and prior to sacrifice. MWM testing was conducted between Days 15-19, followed by PA testing on Days 20-21. At the end of the experiment, rats were anaesthetised with ketamine (90 mg/kg)-xylazine (10 mg/kg) and sacrificed by exsanguination. Brains were bisected into two hemispheres; one hemisphere was used for histological analysis, while the hippocampus from the contralateral hemisphere was used for molecular and biochemical analyses.

The study flow diagram and experimental timeline are shown in Figure 1. The experimental treatment protocol across groups is summarised in Table 1.

**Figure 1.**
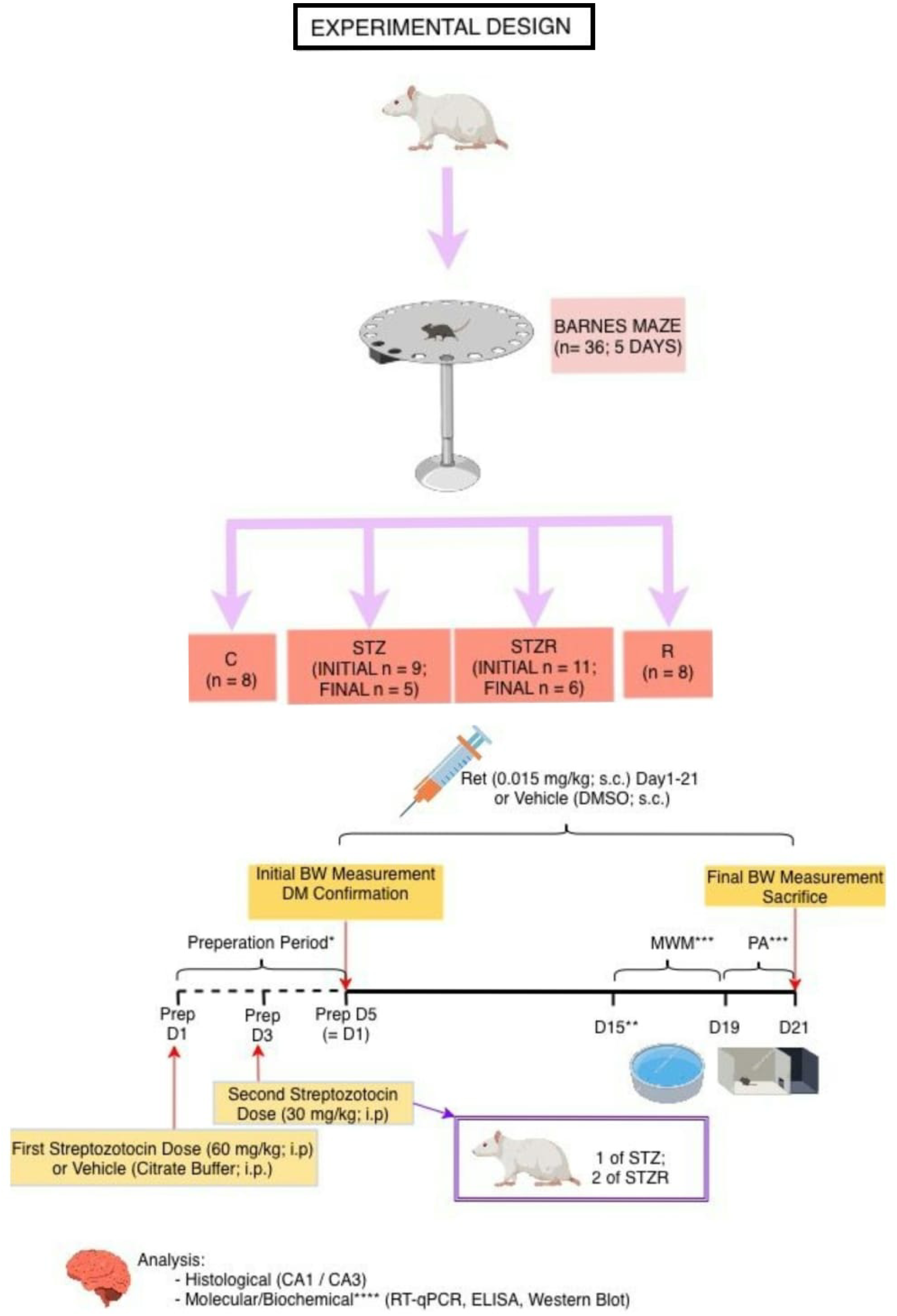
Study flow diagram and experimental timeline. * Blood glucose was assessed on Prep D3 after Streptozotocin administration; animals with glucose levels <250 mg/dL received an additional dose and were re-evaluated on Prep D5 which is also the D1. Treatment was initiated after confirmation of hyperglycaemia. ** Before the behavioural testing, 3 of STZ and 5 of STZR rats did not survive (including the ones which received additional Streptozotocin dose). *** One rat in the STZ group was not included in MWM and PA because of severe clinical deterioration but was still used for histological and molecular analyses. **** Molecular analyses were performed on hippocampal tissue from all rats that survived to the planned sacrifice, including those excluded from behavioural testing because of severe locomotor or clinical deterioration; outliers beyond ±3 SD were removed for IL-1β (1 C, 2 R) and Tau (1 STZR). D: Day; Prep D: Preparation Day; C: Control; STZ: Streptozotocin; STZR: STZ + Retatrutide; R: Retatrutide.

**Table 1.**
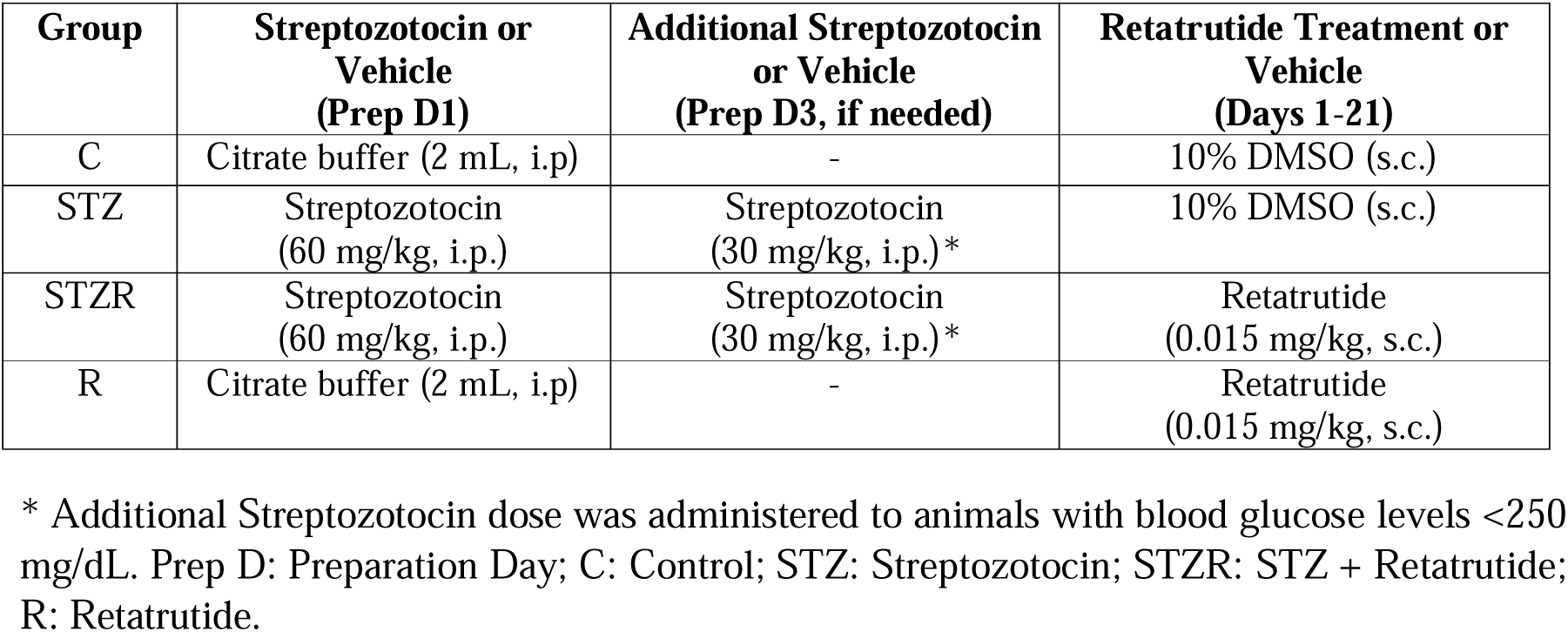
Experimental treatment protocol across study groups.

| Group | Streptozotocin or Vehicle (Prep D1) | Additional Streptozotocin or Vehicle (Prep D3, if needed) | Retatrutide Treatment or Vehicle (Days 1-21) |
| --- | --- | --- | --- |
| C | Citrate buffer (2 mL, i.p) | - | 10% DMSO (s.c.) |
| STZ | Streptozotocin (60 mg/kg, i.p.) | Streptozotocin (30 mg/kg, i.p.)* | 10% DMSO (s.c.) |
| STZR | Streptozotocin (60 mg/kg, i.p.) | Streptozotocin (30 mg/kg, i.p.)* | Retatrutide (0.015 mg/kg, s.c.) |
| R | Citrate buffer (2 mL, i.p) | - | Retatrutide (0.015 mg/kg, s.c.) |

\* Additional Streptozotocin dose was administered to animals with blood glucose levels <250 mg/dL. Prep D: Preparation Day; C: Control; STZ: Streptozotocin; STZR: STZ + Retatrutide; R: Retatrutide.

### 2.5. Behavioural Experiments

#### 2.5.1. Barnes Maze

At the beginning of the study, the Barnes Maze (BM) test was performed to assess spatial learning and to balance baseline performance across groups. The apparatus consisted of a circular platform (122 cm in diameter) with 20 holes (9.5 cm in diameter), one of which led to an escape box. Rats were placed at the centre of the platform for five consecutive days in a random order and allowed 120 s to explore. Animals that located the escape box were kept inside for 15 seconds to reinforce learning [19]. Escape latencies and learning curves obtained over the five days were used to allocate animals into four groups using stratified randomisation.

#### 2.5.2. Morris Water Maze

The Morris Water Maze (MWM) test was conducted in a water tank with a diameter of 130 cm and a depth of 60 cm. The platform was designed in black, and the water in the tank was coloured in black with biosuitable paint to make the platform invisible, and the ambient light was kept dim. The water temperature was maintained at 24 ± 2 °C. The platform’s position did not change during the entire experiment, and cues were placed outside the water tank at visible distances and heights to help the rats learn the location of the hidden platform.

The MWM test was used to assess spatial (hippocampus-dependent) learning and memory. The rats were trained for 5 consecutive days to locate the platform during a maximum of 120 s swimming trials. During each trial, each rat was placed into the tank from one of four directions (north, south, east, or west) in a random order to prevent them from memorising a fixed escape route and to encourage the use of spatial cues. If a rat failed to find the platform within 120 seconds during the first trial, it was gently guided to the platform with the assistance of a stick. After each trial in which the rat located the platform, the rats were allowed to stay on the platform for 30 s to reinforce learning through spatial cues.

The performance of rats over the 5 days was assessed [20] based on their platform-finding time (escape latency) in the MWM. The experiments were recorded using a DMK 22AUC03 monochrome camera. At the end of the experiment, the recorded movement data were analysed using an image recording and processing program developed by faculty members and students at our university’s Faculty of Engineering and Natural Sciences.

#### 2.5.3. Passive Avoidance

The Passive Avoidance (PA) Test was conducted in an apparatus which includes one light and one dark box. While the light box has a steel grating which does not apply electrical shock, the dark box was organised to apply mild electrical shock from its grating. These two boxes were separated by a door which is controlled by the researcher.

During T1 (training session), the rat which placed into the light box is allowed for 60 seconds to explore, and at the 60th second door is opened. As the rat moves into the dark box; door is immediately closed, and a 0.5 mA electrical shock is given for 3 seconds. Following the electrical shock, the session is finalised.

In T2 (1 hour, short-term memory) and T3 (24 hours, medium-long term memory) sessions, the latencies of rats to pass to the dark box from the light box are evaluated. Cut-off value for latencies is determined as 300 seconds [21].

Animals that were unable to complete the MWM and PA procedures according to predefined clinical and locomotor criteria were excluded from behavioural analyses. In contrast, molecular and histological analyses included all animals that survived to the planned sacrifice endpoint. Accordingly, the behavioural and tissue-based datasets were not fully identical in composition. This difference in analysis sets should be considered when interpreting potential discrepancies between behavioural and molecular findings.

### 2.6. Molecular Analyses

Total Tau protein in the hippocampal tissue of sacrificed rats was determined using Western blot, while BDNF, CREB, and AKT gene expression levels were determined through RT-qPCR. Thus, neuroinflammation parameters and cognitive markers were analysed together. Below, the details of the experiment are provided.

#### 2.6.1 RT-qPCR

RT-qPCR experiments were conducted to determine the gene expression levels of BDNF, CREB, and AKT. GAPDH is used as the housekeeping gene. RNA isolation was performed using an RNA isolation kit (Promega, USA) according to the manufacturer’s instructions. RNA concentration and purity were assessed by measuring absorbance at 260 and 280 nm, and RNA integrity was further evaluated by agarose gel electrophoresis. Complementary DNA (cDNA) was synthesised from equal amounts of RNA using a cDNA synthesis kit (Promega, USA).

Quantitative PCR reactions were performed using the Applied Biosystems StepOnePlus Real-Time PCR system, with equal amounts of cDNA loaded for all samples. Melt curve analysis was conducted to verify amplification specificity. Relative gene expression levels were calculated using the ΔΔCt method, and fold changes were determined using the 2^−ΔΔCt formula. For statistical evaluation, ΔCt values were used. All experimental procedures were conducted at the KUYAM laboratory.

#### 2.6.2. Western Blotting Experiments

were conducted to determine Tau protein levels. Tissue samples were separated using Trizol for total protein extraction. The proteins were kept at –80°C until the Western blot procedure. The protein concentration was determined using the bicinchoninic acid (BCA) protein assay method. For the Western blot, the Bio-Rad Mini Protean system (Bio-Rad, USA) was used. Protein samples were separated using a 12% SDS-PAGE gel. Afterwards, they were transferred to a polyvinylidene fluoride (PVDF) membrane and incubated in a blocking buffer. Following this, the samples were incubated overnight at 4°C with the relevant antibodies. After washing steps, the samples were incubated with HRP-conjugated secondary antibodies, and following further washes, the membrane was transferred to a film cassette. Chemiluminescent detection was performed using an ECL kit. The chemiluminescent images were obtained using the Bio-Rad ChemiDoc Image system (Bio-Rad, USA). Band intensities were evaluated densitometrically. All necessary equipment was available at the KUYAM laboratory.

### 2.7. Biochemical Analysis

Inflammation parameters TNF-α and IL-1β were assessed using an ELISA kit.

#### 2.7.1. ELISA Experiments

for measuring TNF-α and IL-1β levels used immunoassay kits following the solid-phase sandwich ELISA (Enzyme-Linked Immuno-Sorbent Assay) principle. The kit procedure was followed, and measurements were taken at the relevant absorbance using a ThermoScan brand plate reader. All necessary equipment was available at the KUYAM laboratory.

### 2.8. Histological Assessment

The brain tissues were immediately placed in a 4% paraformaldehyde solution. The tissues were then embedded in paraffin blocks, from which 4-micron-thick sections were taken. To assess neuronal damage, the CA1 hippocampal region was stained with Haematoxylin & Eosin and Cresyl Violet.

### 2.9. Statistical analysis

Statistical analyses were performed using IBM SPSS Statistics version 26.0 (IBM Corp., Armonk, NY, USA) and GraphPad Prism version 5.01 (GraphPad Software, San Diego, CA, USA). Sample size planning was informed by previous literature using the Streptozotocin model and anticipated effect sizes for behavioural (primarily MWM escape latency) and molecular outcomes, with allowance for expected attrition during the experimental period. Accordingly, an a priori power analysis was conducted using G*Power v3.1 before study initiation, assuming a significance level of 0.05, 80% statistical power, and a large effect size (Cohen’s d ≈ 0.8) for between-group differences in MWM performance, which indicated a planned sample size of approximately 7-8 animals per group.

Normality of data distribution was assessed using the Shapiro-Wilk test. Parametric data are presented as mean ± standard error of the mean (SEM) and were analysed using one-way analysis of variance (ANOVA), followed by Tukey or Games-Howell post hoc tests as appropriate based on the results of Levene’s test for homogeneity of variances. Non-parametric data are expressed as median (interquartile range, IQR) and were analysed using the Kruskal-Wallis test. When a significant overall Kruskal-Wallis effect was detected, post hoc pairwise comparisons were performed using Dunn’s test with Bonferroni-adjusted significance thresholds (α_adj = 0.05 / number of pairwise comparisons) to control for multiple testing.

Performance in the MWM across training days (escape latency and swim speed) was analysed using two-way repeated-measures ANOVA, with Group as the between-subject factor and Day as the within-subject factor. When appropriate, Bonferroni correction was applied for multiple comparisons. Statistical significance was set at p < 0.05 for all analyses. For ELISA and Western blot data, potential outliers were identified using a predefined mean ± 3 SD criterion within each group; values fulfilling these assay-related outlier criteria were excluded from the final analyses.

## 3. Results

### 3.1. Barnes Maze

Prior to group allocation, all animals were tested in the Barnes Maze, and escape latencies across the five baseline training days were used to rank animals by performance. Rats were then assigned to the C, STZ, STZR, and R groups using stratified randomisation such that each group contained animals from comparable performance tiers, thereby ensuring balanced baseline spatial learning (see Supplementary Table S1).

### 3.2. Body weight

The percentage change in body weight was calculated using measurements from Day 1 and just before sacrifice. A one-way ANOVA revealed a significant difference among the groups (*F*(3, 23) = 46.88, *p* < 0.001). Post hoc Games-Howell analysis showed that both diabetic groups (STZ and STZR) exhibited a marked decrease in body weight compared with the control (C) group ((*p* = 0.006 and *p* = 0.003, respectively)) and the Retatrutide-treated (R) group ((*p* = 0.008 and *p* = 0.004, respectively)), with no significant difference between STZ and STZR ((*p* > 0.05)). Mean ± SEM percentage changes in body weight were as follows: C: +5.08 ± 0.49, STZ: −17.71 ± 3.22, STZR: −17.32 ± 3.10, and R: +3.98 ± 0.40. These findings indicate that Retatrutide did not prevent diabetes-induced weight loss, as STZ and STZR showed comparable reductions; in contrast, Retatrutide alone (R) maintained body weight at levels comparable to controls (Fig. 2).

**Figure 2.**
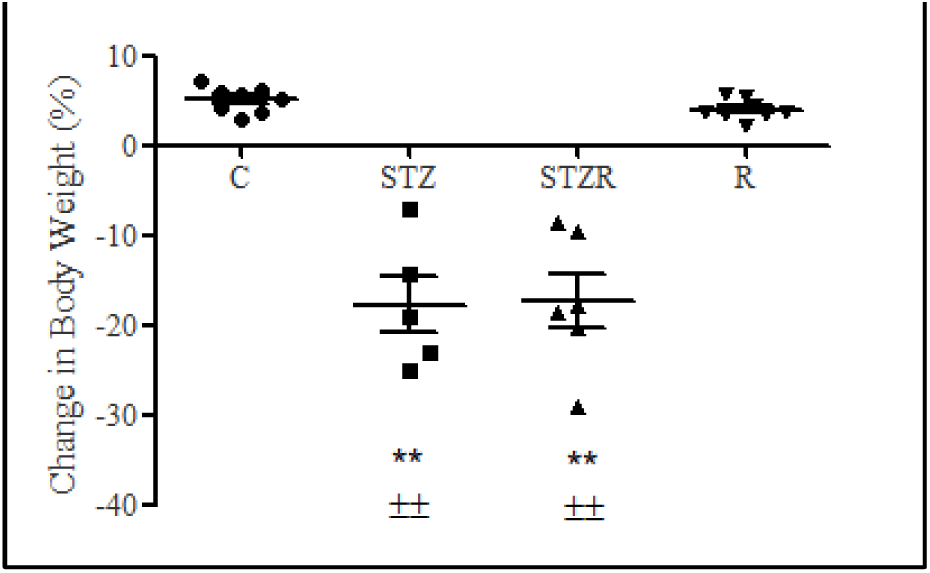
Percentage change in body weight across experimental groups. Data are presented as mean ± SEM (C: n = 8; STZ: n = 5; STZR: n = 6; R: n = 8). Statistical analysis was performed using one-way ANOVA followed by the Games-Howell post hoc test. Symbols indicate significant differences compared with the C (\*\**p* < 0.01), R( ±±*p* < 0.01). C: Control; STZ: Streptozotocin; STZR: STZ + Retatrutide; R: Retatrutide.

### 3.3. Blood glucose

Blood glucose levels were measured on Days 1 and 21 following diabetes induction to assess the development of hyperglycaemia and the effect of Retatrutide treatment (Fig. 3A-B). On Day 1, blood glucose levels differed significantly among the four groups (Kruskal-Wallis *H* = 27.24, *p* < 0.001; Fig. 3A). Post hoc Dunn tests with Bonferroni correction showed that both diabetic groups (STZ and STZR) had significantly higher glucose levels than the control group (C) (both *p* < 0.001), confirming successful diabetes induction at this early time point. No significant difference was detected between the two diabetic groups (STZ vs STZR, *p* > 0.05). Group R displayed glucose levels that did not differ significantly from controls, whereas its values were significantly lower than in both diabetic groups (STZ and STZR) (*p* = 0.021 and *p* = 0.013, respectively), indicating relative protection from Streptozotocin-induced hyperglycaemia.

**Figure 3.**
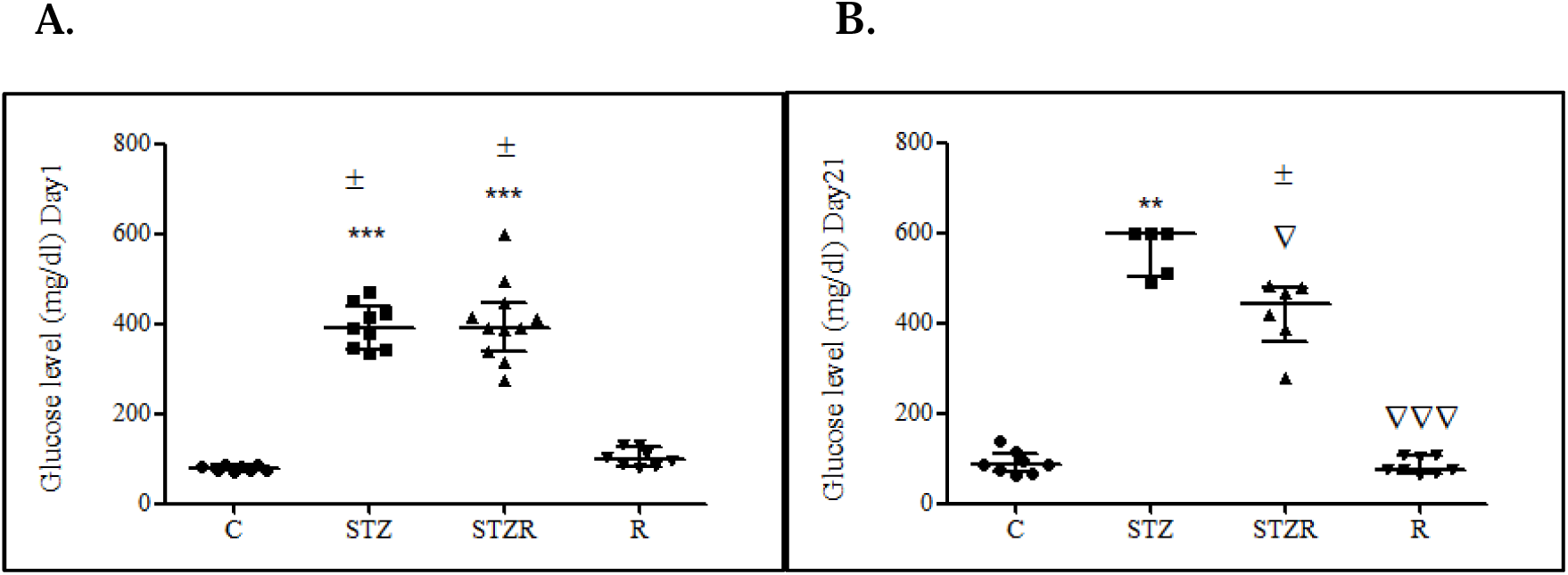
Blood glucose levels on Days 1 and 21 following diabetes induction. (A) Day 1 and (B) Day 21 measurements. Data are presented as median with interquartile range (IQR) (Day 1: C n = 8, STZ n = 9, STZR n = 11, R n = 8; Day 21: C n = 8, STZ n = 5, STZR n = 6, R n = 8). Statistical analysis was performed using the Kruskal-Wallis test followed by Dunn’s post hoc tests for pairwise comparisons with Bonferroni-adjusted significance thresholds. Symbols indicate significant differences compared with the C (\*\**p* < 0.01, \*\*\**p* < 0.001), R (±*p* < 0.05), or STZ group (□*p* < 0.05, □□□*p* < 0.001). C: Control; STZ: Streptozotocin; STZR: STZ + Retatrutide; R: Retatrutide.

By Day 21, group differences in blood glucose remained significant (Kruskal-Wallis *H* = 19.86, *p* < 0.001; Fig. 3B). Dunn post hoc analysis indicated that glucose levels in the STZ group were significantly higher than in the control group (STZ vs C, *p* = 0.003). STZR exhibited significantly lower glucose levels than STZ (*p* = 0.046), supporting an anti-hyperglycaemic effect of Retatrutide. In addition, glucose levels in group R were significantly lower than in the STZ group (*p* = 0.001) and did not differ from those of the control group, suggesting substantial attenuation of Streptozotocin-induced hyperglycaemia, although blood glucose was not fully normalised in all diabetic animals.

### 3.4. Behavioural tests

#### 3.4.1. Morris Water Maze

In the Morris Water Maze (MWM), which evaluates spatial learning based on escape latency, significant differences were observed among the experimental groups. Two-way repeated-measures ANOVA revealed a robust main effect of training day, indicating progressive learning across days (Day: *F*(1.985, 47.628) = 101.34, *p* < 0.001, Greenhouse-Geisser corrected), and a significant main effect of group (Group: *F*(3, 24) = 11.13, *p* < 0.001). Post hoc Bonferroni-corrected pairwise comparisons confirmed that the STZ group exhibited consistently longer escape latencies than the C, STZR, and R groups across the training period (Fig. 4).

**Figure 4.**
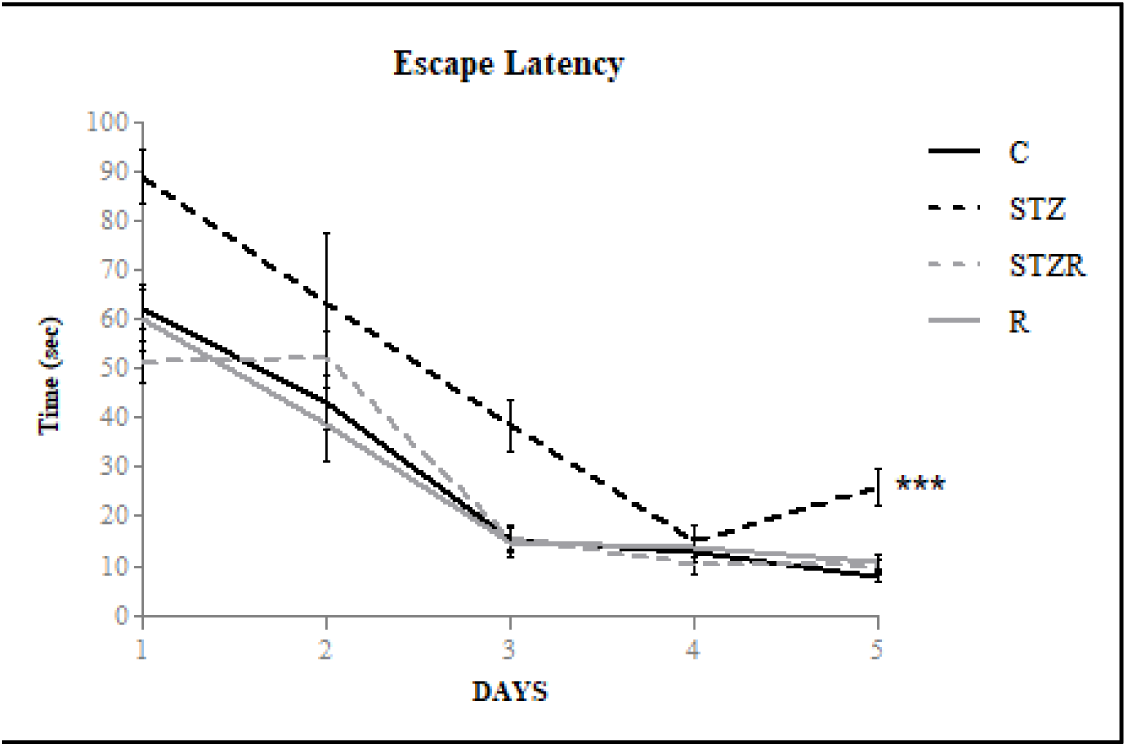
Escape latency in the Morris Water Maze (MWM) across training days. Data are presented as mean ± SEM (C: n = 8; STZ: n = 5; STZR: n = 6; R: n = 8). Statistical analyses were performed using two-way repeated-measures ANOVA with Bonferroni-corrected post hoc comparisons. Symbols indicate statistically significant differences in overall learning curves between the STZ group and the C, STZR, and R groups across training days (\*\*\**p* < 0.001). C: Control; STZ: Streptozotocin; STZR: STZ + Retatrutide; R: Retatrutide.

To further characterise the time course of these differences, exploratory one-way ANOVAs were conducted on individual training days. These analyses indicated significant group effects on Day 1 (*F*(3, 24) = 7.90, *p* = 0.001) and Day 5 (*F*(3, 24) = 15.31, *p* < 0.001), with the STZ group displaying longer escape latencies than all other groups at both time points.

To control for potential locomotor confounding in the interpretation of escape latency, mean swim speed was also analysed across training days using the same repeated-measures framework. A two-way repeated-measures ANOVA with Day as a within-subjects factor and Group as a between-subjects factor showed a significant main effect of Day (Day: *F*(3.329, 76.559) = 8.30, *p* < 0.001, Greenhouse-Geisser corrected), indicating modest adaptation of swimming behaviour over time, but no significant Day × Group interaction (*F*(9.986, 76.559) = 1.42, *p* = 0.25, Greenhouse-Geisser corrected). Importantly, there was no significant main effect of Group on swim speed (Group: *F*(3, 23) = 0.23, *p* = 0.88), demonstrating that average swim speeds were comparable across the C, STZ, STZR, and R groups on all training days (see Supplementary Table S2).

#### 3.4.2. Passive Avoidance

In the PA test, the latency to enter the dark compartment from the illuminated compartment was assessed as an index of memory performance. Because latency data were not normally distributed, non-parametric statistical methods were applied. At the T2 time point (1 h), a significant difference in latency was observed among the groups (Kruskal-Wallis *H* = 9.21, *p* = 0.027; Fig. 5A). Dunn post hoc tests with Bonferroni correction revealed that the STZ group exhibited significantly shorter latencies than the control group (C) (*p* = 0.023), indicating impaired short-term PA memory in untreated diabetic animals. No other pairwise comparisons reached statistical significance at T2, suggesting that the overall group effect at this time point was mainly driven by the contrast between the STZ and control groups.

**Figure 5.**
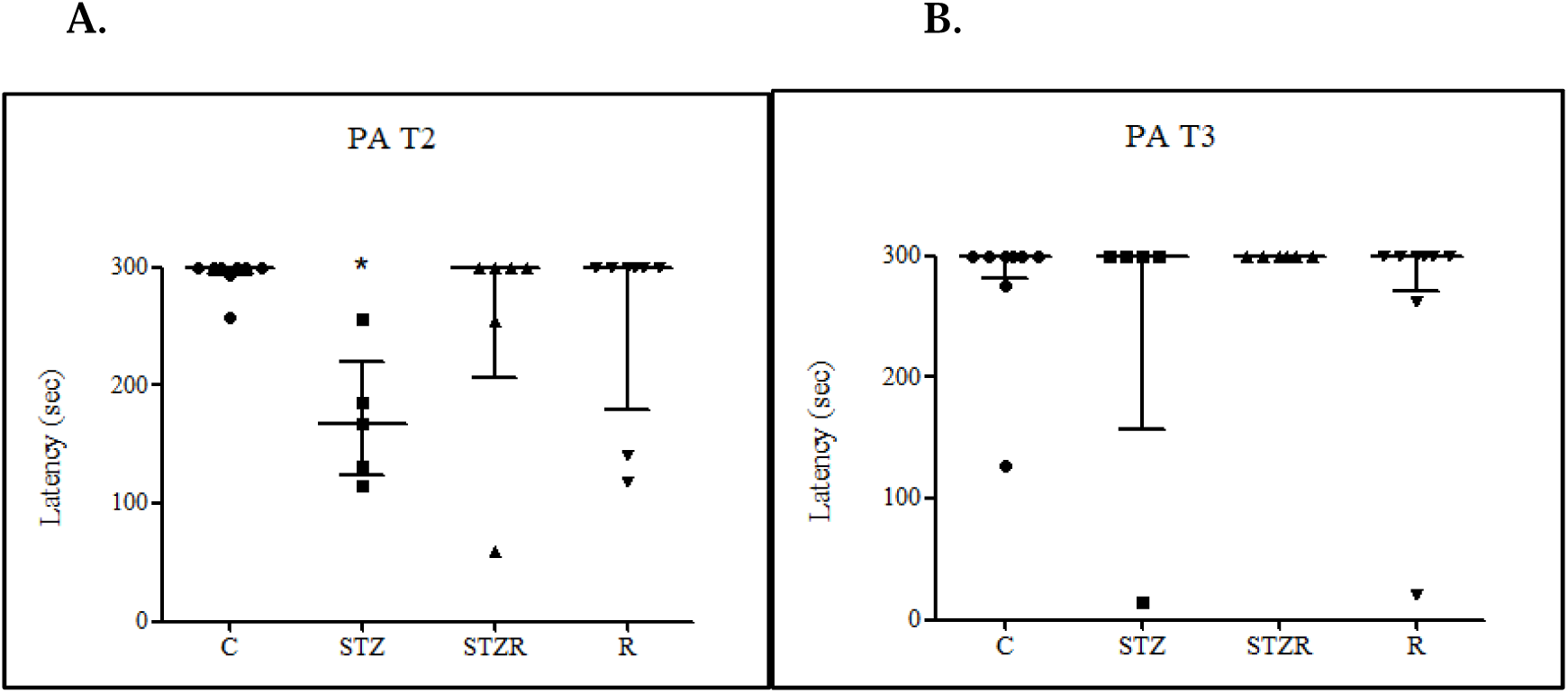
Latency to enter the dark compartment in the passive avoidance (PA) test. (A) Latency measured at T2 (1 h test, short-term memory) and (B) latency measured at T3 (24 h test, long-term memory). Data are presented as median (interquartile range, IQR) (C: n = 8; STZ: n = 5; STZR: n = 6; R: n = 8). Statistical analyses were performed using the Kruskal-Wallis test followed by Dunn’s post hoc tests for pairwise comparisons with Bonferroni-adjusted significance thresholds. Symbols indicate significant differences compared with the C group (* *p* < 0.05). C: Control; STZ: Streptozotocin; STZR: STZ + Retatrutide; R: Retatrutide.

At the T3 time point (24 h), no significant differences in latency were detected among the groups (Kruskal-Wallis *H* = 1.68, *p* = 0.642; Fig. 5B), and therefore no post hoc pairwise tests were performed. Taken together, these findings indicate that short-term, but not long-term, PA performance is selectively impaired in the untreated diabetic group, whereas Retatrutide treatment does not confer a robust, statistically significant protection across groups under the present conditions.

### 3.5. Molecular Analyses

#### 3.5.1. RT-qPCR Experiments

BDNF expression differed significantly among the groups (Kruskal-Wallis *H* = 9.724, *p* = 0.021). Dunn’s post hoc tests with Bonferroni correction showed that BDNF mRNA levels were significantly higher in the R group compared with the control group (C) (*p* = 0.041), whereas no other pairwise comparisons reached statistical significance (Fig. 6A).

**Figure 6.**
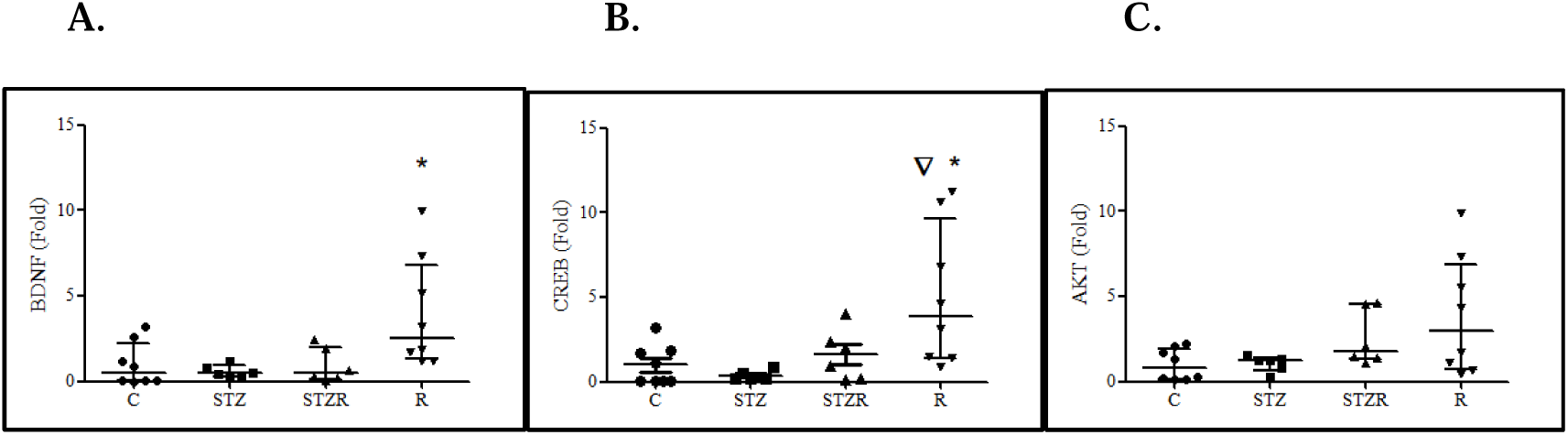
Relative gene expression levels of (A) BDNF, (B) CREB, and (C) AKT determined by RT-qPCR in hippocampal tissue homogenates of experimental groups. Data are presented as median (interquartile range, IQR) (C: n = 8; STZ: n = 6; STZR: n = 6; R: n = 8). Statistical analyses were performed using the Kruskal-Wallis test followed by Dunn’s post hoc tests for pairwise comparisons with Bonferroni-adjusted significance thresholds. Symbols indicate significant differences compared with the C (* *p* < 0.05) or the STZ (□ *p* < 0.05) groups. C: Control; STZ: Streptozotocin; STZR: STZ + Retatrutide; R: Retatrutide.

Similarly, CREB expression differed significantly among the groups (Kruskal-Wallis *H* = 10.573, *p* = 0.014). Post hoc Dunn tests indicated that CREB expression was significantly increased in the R group relative to both the control group (C) (*p* = 0.041) and the STZ group (*p* = 0.028), while the differences between R and STZR and among the remaining group comparisons were not statistically significant (Fig. 6B).

In contrast, AKT mRNA expression did not differ significantly among the experimental groups (Kruskal-Wallis *H* = 5.947, *p* = 0.114; Fig. 6C).

#### 3.5.2. Western Blot Experiments

Western blot analysis of hippocampal homogenates revealed a statistically significant difference in Tau/β-actin levels among the experimental groups (Kruskal-Wallis *H* = 8.065, *p* = 0.045). Dunn’s post hoc tests with Bonferroni correction showed that Tau expression in the control group (C) was significantly higher than in the STZ group (*p* = 0.043), whereas no other pairwise comparisons reached statistical significance (Fig. 7A-B). Thus, streptozotocin-induced diabetes was associated with a reduction in hippocampal Tau protein levels relative to controls, while Retatrutide-treated groups (STZR and R) did not differ significantly from either C or STZ, suggesting only a partial and statistically inconclusive trend toward restoration.

**Figure 7.**
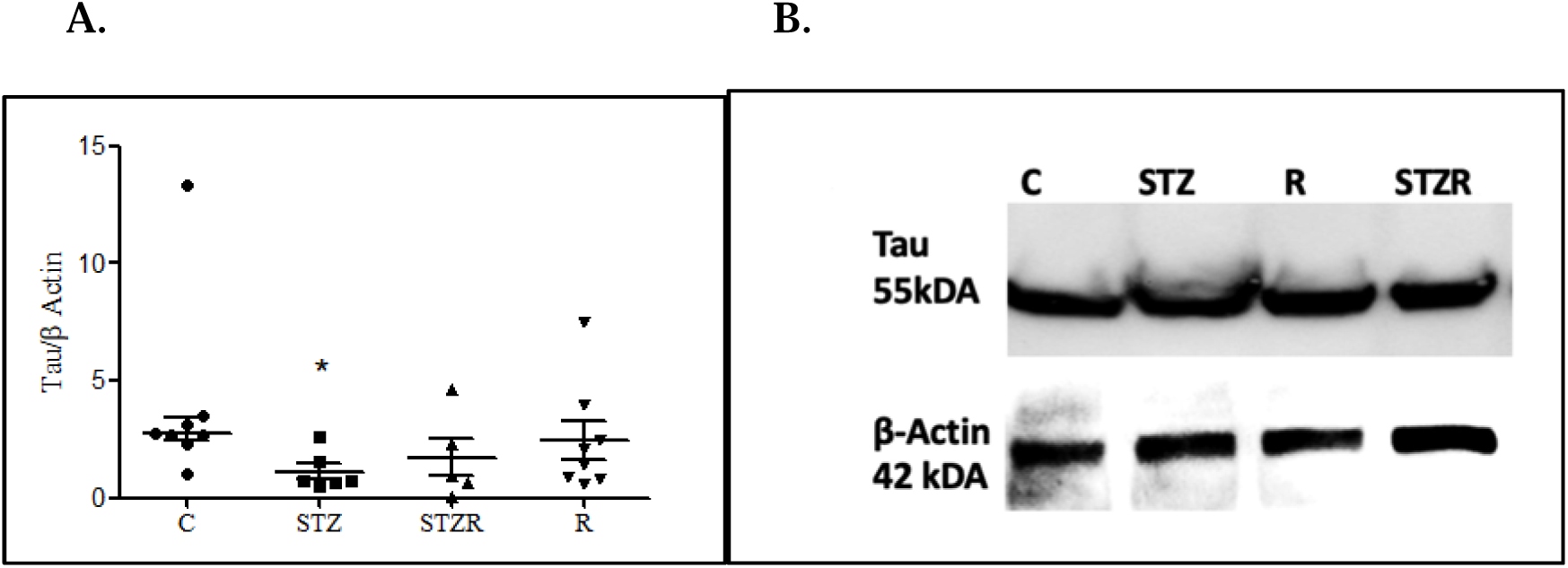
Hippocampal Tau protein expression levels determined by Western blot analysis. (A) Quantitative analysis of Tau/β-actin ratios and (B) representative blot images of Tau (55 kDa) and β-actin (42 kDa) in hippocampal tissue homogenates of experimental groups. Data are presented as median (interquartile range, IQR) (C: n = 8; STZ: n = 6; STZR: n = 5; R: n = 8). Statistical analyses were performed using the Kruskal-Wallis test followed by Dunn’s post hoc tests for pairwise comparisons with Bonferroni-adjusted significance thresholds. Symbols indicate statistically significant differences compared with the C group (* *p* < 0.05). C: Control; STZ: Streptozotocin; STZR: STZ + Retatrutide; R: Retatrutide. In the STZR group, one sample that met predefined assay-related outlier criteria was excluded from the quantitative analysis.

### 3.6. Biochemical Analysis

#### 3.6.1. ELISA Experiments

One-way ANOVA revealed significant differences in IL-1β levels among the experimental groups (** *F*(3, 23) = 6.160, *p* = 0.004**). IL-1β concentrations were expressed as ng/mL, whereas TNF-α concentrations were expressed as mg/L. Post hoc Tukey HSD analysis demonstrated that IL-1β concentrations were significantly higher in the STZ group compared to the control group (C) (*p* = 0.002) and the R group (*p* = 0.030), whereas no significant difference was observed between the STZ and STZR groups (*p* = 0.137) (Fig. 8A). TNF-α levels also differed significantly among the experimental groups (** *F*(3, 23) = 3.622, *p* = 0.027**). Post hoc Tukey HSD analysis indicated that TNF-α concentrations were significantly higher in the STZ group compared to the STZR group (*p* = 0.030), whereas no statistically significant difference was detected between the STZ and control groups (C) (*p* = 0.062) or between the STZ and R groups (*p* = 0.089) (Fig. 8B).

**Figure 8.**
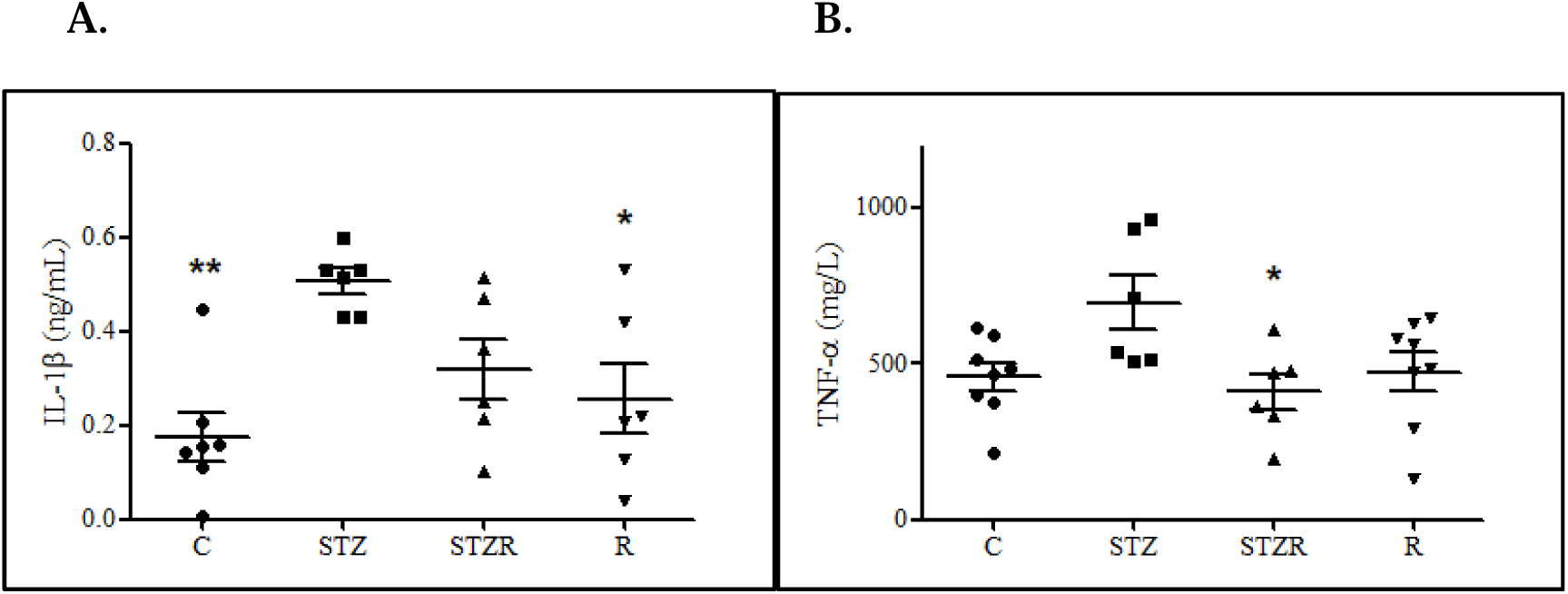
Hippocampal cytokine levels measured by ELISA. (A) IL-1β and (B) TNF-α concentrations in hippocampal tissue homogenates of experimental groups. Data are presented as mean ± SEM (IL-1β: C: n = 7; STZ: n = 6; STZR: n = 6; R: n = 6; TNF-α: C: n = 8; STZ: n = 6; STZR: n = 6; R: n = 8). Statistical analyses were performed using one-way ANOVA followed by Tukey’s post hoc test. Symbols indicate statistically significant differences relative to the STZ group within each panel (* *p* < 0.05, ** *p* < 0.01). C: Control; STZ: Streptozotocin; STZR: STZ + Retatrutide; R: Retatrutide. For IL-1β, one value in the C group and two values in the R group that fulfilled predefined assay-related outlier criteria were excluded from the final analysis to avoid undue distortion of variance estimates.

### 3.7. Histological Assessment

#### 3.7.1. Cortical Evaluation

A normal histological architecture was observed in the cerebral cortex of the C group, with preserved cortical layers and neurons displaying normal morphology. In the STZ group, neuronal density appeared reduced, and many neurons were surrounded by prominent vacuoles, accompanied by evident dilatation of perivascular spaces. In the STZR group, neuronal density was relatively increased compared to the STZ group, and the number of vacuoles surrounding neurons was decreased; however, perivascular dilatation remained apparent, similar to that observed in the STZ group. The R group displayed cortical morphology largely comparable to that of the C group, with minimal vacuolisation and preserved perivascular architecture.

#### 3.7.2. Hippocampal Evaluation

In the hippocampus, the C group exhibited normal histological architecture in the CA1 and CA3 regions, with well-organised pyramidal cell layers and neurons showing normal nuclear and cytoplasmic morphology. In the STZ group, both CA1 and CA3 regions showed shrunken pyramidal neurons with pyknotic nuclei, cytoplasmic vacuolisation, and disorganisation of the pyramidal cell layer, consistent with diabetic neurodegeneration. In the STZR group, the CA1 region largely retained a normal histological appearance, whereas in CA3 mild disorganisation of the pyramidal layer and occasional shrunken neurons with pyknotic nuclei were still observed. The R group demonstrated preserved CA1 architecture similar to controls, while neuronal arrangement in the CA3 region was mildly disorganised but comparable to that observed in the STZR group, suggesting partial structural preservation following Retatrutide treatment.

## 4. Discussion

In this study, the effects of Retatrutide on metabolic parameters and cognitive performance in diabetic and non-diabetic rats were evaluated from both physiological and behavioural perspectives in an insulin□deficient, Streptozotocin□induced diabetes model (present study). The findings demonstrated that Streptozotocin-induced diabetes was associated with marked metabolic disturbances and cognitive impairment, whereas Retatrutide treatment was associated with a partial attenuation of these alterations rather than complete normalisation (present study). As a novel GIP/GLP□1/Glucagon triple receptor agonist, Retatrutide has recently gained significant attention for its superior efficacy in metabolic disorders; however, its potential to counteract diabetes□associated cognitive decline represents an emerging frontier in neuropharmacology [22]. The insulin-deficient Streptozotocin model (T1DM-like) was selected to investigate diabetes-induced neurocognitive impairment, with an emphasis on mechanisms driven by hyperglycaemia and neuroinflammation rather than on a specific diabetes subtype [23,24]. Accordingly, the present findings should be interpreted within the framework of severe insulin□deficiency-driven metabolic and neuroinflammatory stress, rather than as a direct model of type 2 diabetes.

The pronounced body weight loss and persistent hyperglycaemia observed in the STZ group are consistent with the severe insulin deficiency and catabolic state characteristic of this model. Retatrutide treatment did not prevent body weight loss, which may be explained by the dominant insulin deficiency and enhanced lipolysis and proteolysis masking potential effects on weight regulation. Although GLP-1/triple agonists show metabolic benefits [4,6], their effects on weight loss remain limited in insulin-deficient Streptozotocin models [25].

In contrast, detailed blood glucose analyses showed that Retatrutide exerted a robust anti□hyperglycaemic effect, particularly in non□diabetic rats. On Day 1, both diabetic groups (STZ and STZR) displayed the expected marked hyperglycaemia compared with controls, whereas Retatrutide□treated non□diabetic animals (R) had glucose levels comparable to controls but significantly lower than those of both diabetic groups, indicating early protection against Streptozotocin□induced glycaemic elevation. By Day 21, hyperglycaemia persisted in the STZ group, while Retatrutide□treated diabetic rats (STZR) and non□diabetic rats (R) exhibited significantly lower glucose levels than STZ animals, with R values indistinguishable from controls. These findings suggest that Retatrutide markedly attenuates Streptozotocin□induced hyperglycaemia, bringing glycaemia close to the control range in non□diabetic animals and partially improving glycaemic control in diabetic rats, despite ongoing β□cell loss [17].

Behavioural assessments indicated that Streptozotocin-induced diabetes was associated with marked impairments in task performance, whereas Retatrutide treatment was associated with a partial attenuation of these deficits rather than a complete normalisation. In the Morris Water Maze (MWM), spatial learning was markedly disrupted in the STZ group, as reflected by prolonged escape latencies, while the performance of the STZR and R groups remained comparable to that of controls (Fig. 4).The main group effect across training days was robust, and the absence of a significant difference between the C and STZ groups on Day 4, despite a clear separation on Day 5, likely reflects variability at an intermediate stage of learning when control and Retatrutide-treated animals were still approaching asymptotic performance, whereas STZ rats remained lagging behind; this pattern is compatible with a gradual divergence in spatial learning curves rather than with an abrupt change in locomotor capacity (Fig. 4). In line with this interpretation, mean swim speed did not differ significantly between groups across training days, arguing against a dominant contribution of gross locomotor slowing to the latency differences and supporting an interpretation primarily in terms of diabetes-related disruption of spatial learning rather than of non-specific motor impairment (see Supplementary Table S2). Nevertheless, because escape latency is inherently influenced by both cognitive and motor factors, the present findings are best interpreted as evidence that Retatrutide preserves overall MWM performance rather than as definitive proof of completely intact reference memory.

In the Passive Avoidance (PA) task, the updated analyses confirmed that short□term avoidance performance at the T2 time point was selectively impaired in the STZ group: STZ rats showed significantly shorter latencies than controls, whereas neither the STZR nor the R groups differed significantly from the control group, which is compatible with a partial prevention of the Streptozotocin□associated deficit in this task rather than a fully normalised profile. At the T3 time point (24 h), between-group differences were no longer statistically significant, although STZ animals tended to show numerically shorter latencies, suggesting a non□significant trend toward weakened long-term memory retention (Fig. 5). Because the PA task was administered after completion of MWM training, the T2 assessment probed short□term aversive memory on the background of accumulated task exposure and stress; under these conditions, the selective deficit in the STZ group, together with preserved performance in STZR and R rats, suggests an early diabetes□related vulnerability that may be at least partly counteracted by Retatrutide under the present experimental conditions. Taken together, these findings indicate that diabetes particularly compromises short□term avoidance□related memory, and that Retatrutide may confer limited but detectable protection against this early deficit without producing a clearly superior long□term retention profile across groups. The combined pattern across MWM and PA therefore suggests that Streptozotocin□induced diabetes most strongly affects tasks requiring sustained spatial learning and short□term aversive memory, whereas Retatrutide preserves overall behavioural performance across both paradigms, albeit with task□ and time□dependent variation in effect size.

Overall, the combined MWM and PA results indicate that diabetes adversely affects learning-and memory-related task performance, while Retatrutide is associated with an improvement and partial preservation of behavioural performance rather than a complete restoration to control levels. Importantly, the inclusion of swim-speed analyses indicates that group differences in MWM escape latency are unlikely to be driven by gross locomotor slowing, although more detailed locomotor control indices (such as path length, thigmotaxis, and probe-trial search patterns) would be required to fully disentangle mnemonic from non-mnemonic components. These findings are consistent with previous studies reporting potential cognitive benefits of GLP-1-related and multi-incretin agonists [7,26–31]. Triple-receptor agonism demonstrates superior neuroprotection over mono-therapy, likely through synergistic GIP/GLP-1/Glucagon effects on synaptic plasticity [32].

Cognitive improvement cannot be attributed solely to metabolic recovery, as the dissociation between glycaemic normalisation and persistent body weight loss, together with preserved swim speed despite impaired latency performance, suggests additional central or peripheral-central mechanisms beyond pure glucose control. Recent pharmacokinetic insights suggest that multi-incretin agonists can cross the blood-brain barrier and directly modulate neuronal receptors, providing neuroprotection independent of peripheral glucose levels [33]. However, blood-brain barrier penetration has not yet been demonstrated specifically for Retatrutide after subcutaneous administration. Therefore, the behavioural and molecular effects observed in the present study should be interpreted as being consistent with either direct central actions or indirect peripheral-central signalling, rather than as definitive evidence that s.c. Retatrutide reaches the brain parenchyma.

Suppression of neuroinflammation appears to be a key component underlying the cognitive effects of Retatrutide. The tendency toward reduced IL-1β and TNF-α levels in the STZR group, compared with the marked elevations observed in the STZ group, with TNF-α reduction reaching statistical significance, supports the role of neuroinflammation in diabetes-related cognitive impairment. IL-1β is known to disrupt long-term potentiation and synaptic transmission [34] while TNF-α alters postsynaptic AMPA receptor regulation [35], thereby impairing synaptic homeostasis. Previous studies reporting that GLP-1-based agents suppress microglial activation, inhibit NF-κB signalling, and reduce mitochondrial stress [36], further support the notion that Retatrutide may alleviate inflammatory burden, particularly through modulation of TNF-α. Recent research demonstrates that Retatrutide (a triple agonist) significantly suppresses microglial nitrite production, TNF-α release, and COX2 expression in LPS-activated microglia, outperforming single GLP-1R agonists like exendin-4 [37].

Cellular survival pathways closely linked to neuroinflammation also appear to be affected in the diabetic brain. Although no statistically significant differences were detected in AKT mRNA expression among groups, the downward trend observed in the STZ group may reflect impairment within the IRS-1/PI3K/AKT signalling axis, which has been associated with GSK-3β activation under diabetic conditions [38]. The modest upward trend in AKT mRNA expression in the STZR group suggests partial preservation of transcriptional capacity; however, functional activation of AKT signalling is regulated primarily at the post-translational level and cannot be inferred from the present data, which are limited to mRNA expression.

When inflammation-related cellular dysfunction is considered, the revised RT-qPCR data further highlight alterations in neurotrophic signalling pathways, in line with experimental work showing that GLP□1 receptor activation enhances cAMP/PKA□dependent CREB signalling and BDNF expression in the diabetic and degenerating brain [30,39]. Although BDNF and CREB expression levels in the STZ group did not differ significantly from controls, reduced median values are compatible with a relative suppression of neurotrophic support in the diabetic brain [30]. Importantly, Retatrutide monotherapy in non-diabetic rats (R group) produced a significant up-regulation of both BDNF and CREB at the mRNA level, while Retatrutide-treated diabetic rats (STZR) showed an intermediate pattern without significant changes relative to either C or STZ.This profile suggests that Retatrutide can actively modulate BDNF-CREB signalling at the transcriptional level under non□diabetic conditions and may partially counteract diabetes□related vulnerability, even when clear normalisation is not achieved, consistent with clinical evidence that GLP□1 receptor agonists improve cognitive outcomes in patients with type 2 diabetes [31] and with preclinical data demonstrating that GLP[1[based and multi-receptor agonists, including tirzepatide, strengthen the BDNF-TrkB-CREB axis and protect against neurodegeneration [39,40].

Consistent with these molecular findings, the updated Western blot analyses indicate that structural alterations related to neuronal integrity are already evident at the level of total Tau. Hippocampal total Tau/β-actin ratios were significantly reduced in STZ animals compared with controls, whereas neither STZR nor R groups differed significantly from either C or STZ. The significant reduction in total Tau levels observed in the STZ group reflects compromised microtubule stability and extensive neuronal loss rather than pathological Tau hyperphosphorylation[41]. Since Tau is a critical microtubule-associated protein stabilising the neuronal cytoskeleton, its depletion indicates severe cytoskeletal destabilisation and axonal degeneration in the diabetic brain [42]. 2024 literature emphasises that in chronic diabetic states, total Tau depletion serves as a definitive biomarker of advanced neuronal loss and cytoskeletal collapse [25]. In the present study, Retatrutide-treated groups showed Tau levels that were numerically higher than those of STZ rats but not statistically distinct, indicating at most a partial and statistically inconclusive trend toward structural preservation (Fig. 7). These molecular observations were corroborated by histopathological examination, which provided converging anatomical evidence of attenuation of diabetes-related neurodegeneration (Fig. 9). The STZ group exhibited characteristic features of diabetic encephalopathy, including severe neuronal degeneration, pyknotic nuclei, cytoplasmic vacuolization, and disorganisation of the pyramidal cell layers in the hippocampus and cerebral cortex [43]. These morphological changes are often attributed to chronic hyperglycemia-induced oxidative stress and neuroinflammation. Conversely, Retatrutide administration visibly attenuated these degenerative features. The STZR group displayed a relatively preserved cytoarchitecture with reduced neuronal loss and fewer signs of vacuolization compared to untreated diabetic rats (Fig. 9). This structural preservation implies that the anti-inflammatory actions of Retatrutide may contribute to maintaining the neural circuits required for cognitive function.

**Figure 9.**
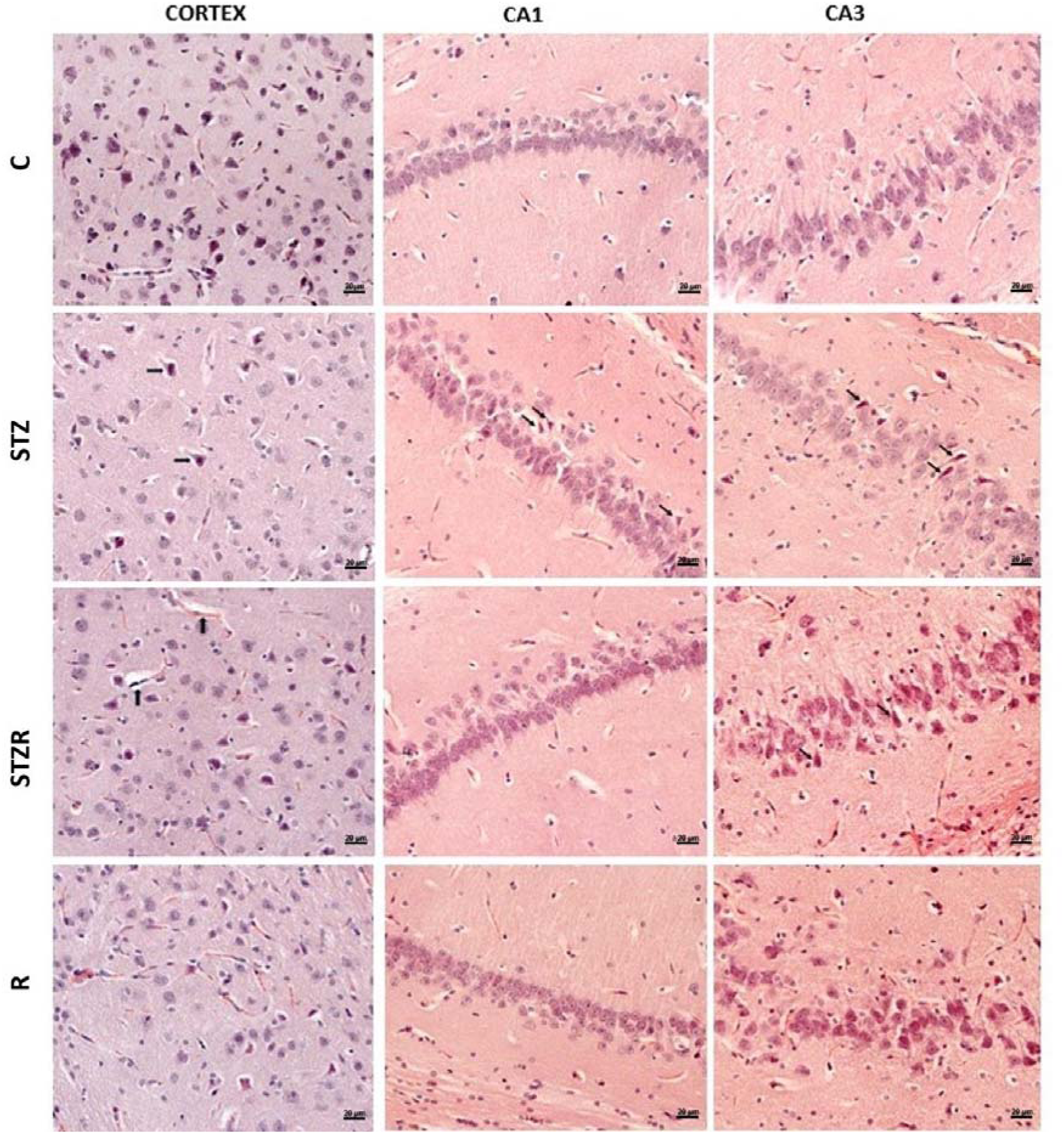
Representative histological images of the cerebral cortex and hippocampus following experimental treatments. Hematoxylin and eosin (H&E)-stained sections show the cerebral cortex and hippocampal CA1 and CA3 regions in the control (C), Streptozotocin (STZ), Streptozotocin + Retatrutide (STZR), and Retatrutide (R) groups. Cortical sections were additionally stained with Cresyl violet. Arrows indicate key histopathological features observed across groups: right-pointing arrows denote vacuolated neurons, upward-pointing arrows indicate perivascular space enlargement, and cross-shaped arrows indicate shrunken pyramidal neurons with pyknotic nuclei. Scale bar: 20 µm. C: Control; STZ: Streptozotocin; STZR: STZ + Retatrutide; R: Retatrutide.

Taken together, Retatrutide’s cognitive and molecular effects in this model should be interpreted as suggesting a multifaceted, but not uniform, attenuation of diabetes□related brain alterations beyond glycaemic control, rather than as definitive evidence of a clear and uniform protective effect across all behavioural and molecular domains. Neuroinflammation suppression, transcriptional preservation of BDNF-CREB signalling, and hippocampal cytoarchitecture maintenance emerge as convergent mechanisms that may underlie the observed improvements. Another limitation of the present study is that RT□qPCR measurements of BDNF, CREB, and AKT were not complemented by protein□level assays for these targets; therefore, our conclusions regarding neurotrophic and survival□related signalling are based on transcriptional evidence only and warrant confirmation by Western blot or proteomic approaches in future work.

The Streptozotocin paradigm was used here as a chemically induced diabetes model suitable for mechanistic evaluation of behavioural and hippocampal molecular alterations. However, this model does not capture the full clinical and pathophysiological heterogeneity of human diabetes, and the present findings should therefore be interpreted primarily within a controlled preclinical framework.

Importantly, the Streptozotocin regimen used here induces an insulin□deficient, T1DM□like phenotype rather than a classical type 2 diabetes model, so our results should be viewed as reflecting Streptozotocin□induced diabetes□related neurocognitive impairment rather than the full clinical and pathophysiological heterogeneity of type 2 diabetes mellitus, and the present findings should therefore be interpreted primarily within a controlled preclinical framework.

Retatrutide effects were evaluated over a limited treatment period, in male rats only, and within a restricted set of molecular pathways, and histopathological assessment was primarily qualitative. In addition, behavioural outcomes relied mainly on escape latency and avoidance latency measures; although swim speed was comparable between groups, we did not include probe trials, path-length quantification, or detailed analyses of search strategies, which would provide more fine-grained locomotor and mnemonic control indices and help to more clearly dissociate cognitive performance from potential subtle motor or motivational influences. Finally, the sample sizes in some experimental readouts were modest and required non-parametric statistics, which, despite appropriate multiple-comparison control, may limit sensitivity to small effects. Therefore, these findings should be validated in models more closely resembling type 2 diabetes mellitus, extended to both sexes, and complemented by comprehensive molecular, functional, and behavioural assessments in future studies.

